# Commonly used Hardy-Weinberg equilibrium filtering schemes impact population structure inferences using RADseq data

**DOI:** 10.1101/2021.06.15.448615

**Authors:** William S. Pearman, Lara Urban, Alana Alexander

## Abstract

Reduced representation sequencing (RRS) is a widely used method to assay the diversity of genetic loci across the genome of an organism. The dominant class of RRS approaches assay loci associated with restriction sites within the genome (restriction site associated DNA sequencing, or RADseq). RADseq is frequently applied to non-model organisms since it enables population genetic studies without relying on well-characterized reference genomes. However, RADseq requires the use of many bioinformatic filters to ensure the quality of genotyping calls. These filters can have direct impacts on population genetic inference, and therefore require careful consideration. One widely used filtering approach is the removal of loci which do not conform to expectations of Hardy-Weinberg equilibrium (HWE). Despite being widely used, we show that this filtering approach is rarely described in sufficient detail to enable replication. Furthermore, through analyses of *in silico* and empirical datasets we show that some of the most widely used HWE filtering approaches dramatically impact inference of population structure. In particular, the removal of loci exhibiting departures from HWE after pooling across samples significantly reduces the degree of inferred population structure within a dataset (despite this approach being widely used). Based on these results, we provide recommendations for best practice regarding the implementation of HWE filtering for RADseq datasets.

## Introduction

Reduced representation sequencing (RRS) is a population genomic approach that enables assaying of a reduced set of genetic loci across the genome of an organism. There are many reduced representation sequencing approaches, some of which assay loci associated with restriction sites within the genome, including approaches such as Genotyping-by-Sequencing (GBS), Restriction site-Associated DNA sequencing (RADseq), double digest RADseq (ddRADseq), DArTSeq, and hybridization of RAD probes (hyRAD) (see (Andrews et al., 2016) for a discussion and summary of these methods). These approaches are an efficient and, in comparison with Whole-Genome Sequencing (WGS), cost-efficient method for generating population genomic datasets, often with a focus on inferring population structure of non-model organisms. The uniting feature of these different approaches is utilising restriction sites in an attempt to assess genome-wide diversity while not having to sequence the complete genome. For the remainder of this paper, we group these various approaches under the umbrella term of “RADseq”.

The application of RADseq, particularly to non-model organisms, however, can pose particular challenges. First, RADseq can be affected by allelic dropout, the failure to identify an allele due to the loss of a restriction site which leads to missing data for that allele and therefore an apparent reduction in heterozygosity in samples (Cooke et al., 2016). Furthermore, the inferences drawn from RADseq data originating from non-model species often depend on the availability of a reference genome of the species of interest or a closely related one (Galla et al., 2019). While a reference genome is not essential for conducting analyses based on RADseq datasets, *de novo* assembly without a reference can result in more misassembled genetic loci (LaCava et al., 2020). However, as RADseq typically produces a large amount of data, bioinformatic filtering approaches can be leveraged to adjust for the potential biases of RADseq approaches.

The application of such filters help to normalize RADseq data across experiments, and to check if the data is consistent with the assumptions made by downstream analyses (O’Leary et al., 2018). For population structure inference in non-model species (Choquet et al., 2019), downstream analyses often make assumptions about factors such as the population size (i.e. very large), the sampling scheme (i.e. randomized sampling), and the species in question (i.e. diploid). Ordination techniques such as Principal Component Analysis (PCA) are therefore often used for preliminary analysis of RADseq data since they do not rely on these assumptions, however, they lack the translation to population parameters that parametric approaches such as admixture analyses or F-statistics offer (Falush et al., 2003; Wright, 1943).

One commonly used admixture approach is STRUCTURE, a widely used tool for identifying distinct genetic groups in population genetic data, and for subsequently analysing the degree of admixture between individuals (Falush et al., 2003; Porras-Hurtado et al., 2013). STRUCTURE iteratively clusters individuals into groups in order to minimise the Hardy-Weinberg disequilibrium (HWD) within groups while maximising it between groups (Pritchard et al., 2010). Thus, STRUCTURE makes explicit assumptions about the relationship between HWD and genetic structure within groups.

F-statistics are frequently used to infer the degree of genetic structure within predefined groups based on observed heterozygosity relative to expected heterozygosity. Population structure is typically measured using F_ST_, which is defined as the relative reduction in heterozygosity due to partitioning the total dataset into putative populations (Whitlock, 2011; Wright, 1943). Accurate *a priori* delineation of groups or ‘populations’ is essential for leveraging F_ST_ to characterise population structure (De Meeûs, 2018). F_ST_ can further be influenced by independent factors that impact the heterozygosity of individual SNPs (Single Nucleotide Polymorphisms) (such as natural selection or technological artifacts including null alleles; De Meeûs, 2018; Meirmans & Hedrick, 2011; Whitlock, 2011).

The assumptions of the various methods highlighted here reinforce the need for appropriate bioinformatic filtering approaches when inferring population structure from RADseq data. Filtering approaches can substantially influence the inference of genetic structure, especially when filters disproportionately affect potentially informative loci (Graham et al., 2020; Shafer et al., 2017). Linck & Battey (2019) showed that minor allele frequency (MAF) filtering of datasets may be problematic since it alters the site frequency spectrum (SFS) across loci according to their rate of missingness. Additional recent work has revealed that both variant call rate and MAF can affect population genetic inferences and genotype-environment association studies (Ahrens et al., 2021; Selechnik et al., 2020). In Table 1, we summarise filtering approaches that are commonly applied to RADseq data, the reasons for their usage, and how they can affect population genetic inference.

**Table 1.** Description of commonly used filtering approaches in the analysis of RADseq data (“Filter”), the reason for their usage (“Usage”), and how they impact population genomic inference (“Impact”).

| Filter | Usage | Impact | Reference |
| --- | --- | --- | --- |
| Hardy-Weinberg equilibrium (HWE) | Removes loci under selection<br>Removes library and sequencing artifacts | **Unknown** | (Gruber et al., 2018; Sethuraman et al., 2019; Waples, 2015) |
| Linkage within loci | Mitigates effects of non-independence of Single Nucleotide Polymorphisms (SNPs) by removing physically linked SNPs. | Reduces false signals of population structure<br>Necessary for STRUCTURE (If LD correction is not used) | (O’Leary et al., 2018) |
| Locus level diversity | Loci with high SNP density (i.e. many SNPs within a locus) may be the result of polyploidy | Can remove putative paralogous loci | (Hohenlohe et al., 2011; Mastretta-Yanes et al., 2015) |
| Minor Allele Frequency (MAF)/Count (MAC) | Identification of genotyping errors | Can remove informative loci if not applied carefully<br>MAF will affect loci differently based on missingness<br>Removes genotyping errors | (Linck & Battey, 2019; O’Leary et al., 2018) |
| Variant call rate | Ensures SNP panel is well represented across individuals | Can dramatically reduce number of loci | (O’Leary et al., 2018) |
|  |  | • Helps ensure samples are comparable |  |

The removal of genetic loci exhibiting departures from Hardy-Weinberg Equilibrium (HWE) is a commonly applied filter (Waples, 2015). HWE describes the state of an ideal population in the absence of evolutionary forces, where allele frequencies are predictable since they remain constant across generations (Garnier‐Géré & Chikhi, 2013). The removal of genetic loci departing from HWE is often used to remove genotyping errors (Hendricks et al., 2018) and loci that are potentially under selection (Lachance, 2009; Wang et al., 2005). The removal of genotyping errors is, in general, beneficial for downstream analyses, while the removal of loci under selection may be required for analyses that assume neutrality of loci. However, many other factors can cause departures from HWE, especially since the assumptions of HWE are rarely met in real biological populations (Waples, 2015), and therefore the removal of loci out of HWE may have substantial effects on population genetic inferences.

The, arguably, most obvious other factor that can cause departures from HWE is the Wahlund effect, where heterozygosity is dramatically reduced due to the inadvertent pooling of multiple populations (De Meeûs, 2018). Excessive deviation from HWE heterozygosity expectations can also arise from repetitive genomic elements (Hohenlohe et al., 2011). Other scenarios that lead to HWE departure that are also frequently observed in real populations include overlapping generations, non-panmictic reproduction, non-diploidy, and very small population sizes. Genotype/SNP (Single Nucleotide Polymorphism) calling approaches represent further potential sources of departure from HWE: Genotype calling can be sensitive to sequencing depth, and to the number of mismatches allowed to call a variant, both of which can lead to a reduction in heterozygosity and in turn lead to HWE departures (Cumer et al., 2021).

While the impact of such factors is often minor, genetic inferences for species which have many potential causes of HWE departures (such as endangered species) might be heavily impacted by decisions around HWE-based filtering. Specifically, when conservation decisions are based on genetic inferences that utilize HWE filtering, it is essential to ensure that this is done appropriately to aid in the management of already vulnerable populations.

The question of if and how a genomic dataset should be filtered for departure from HWE is a difficult one. Sample stratification has to be taken into account; genetic loci that depart from HWE can be filtered in various ways (Fig. 1): No loci removed based on HWE departures (‘No Filter’), loci removed if they exhibit departures in any sampling location (‘Out Any’), loci removed if they exhibit departures from HWE in all sampling locations (or a certain proportion of sampling locations) (‘Out All’, ‘Out Some’), or loci removed if they exhibit departures across sampling locations (‘Out Across’).

**Figure 1.**
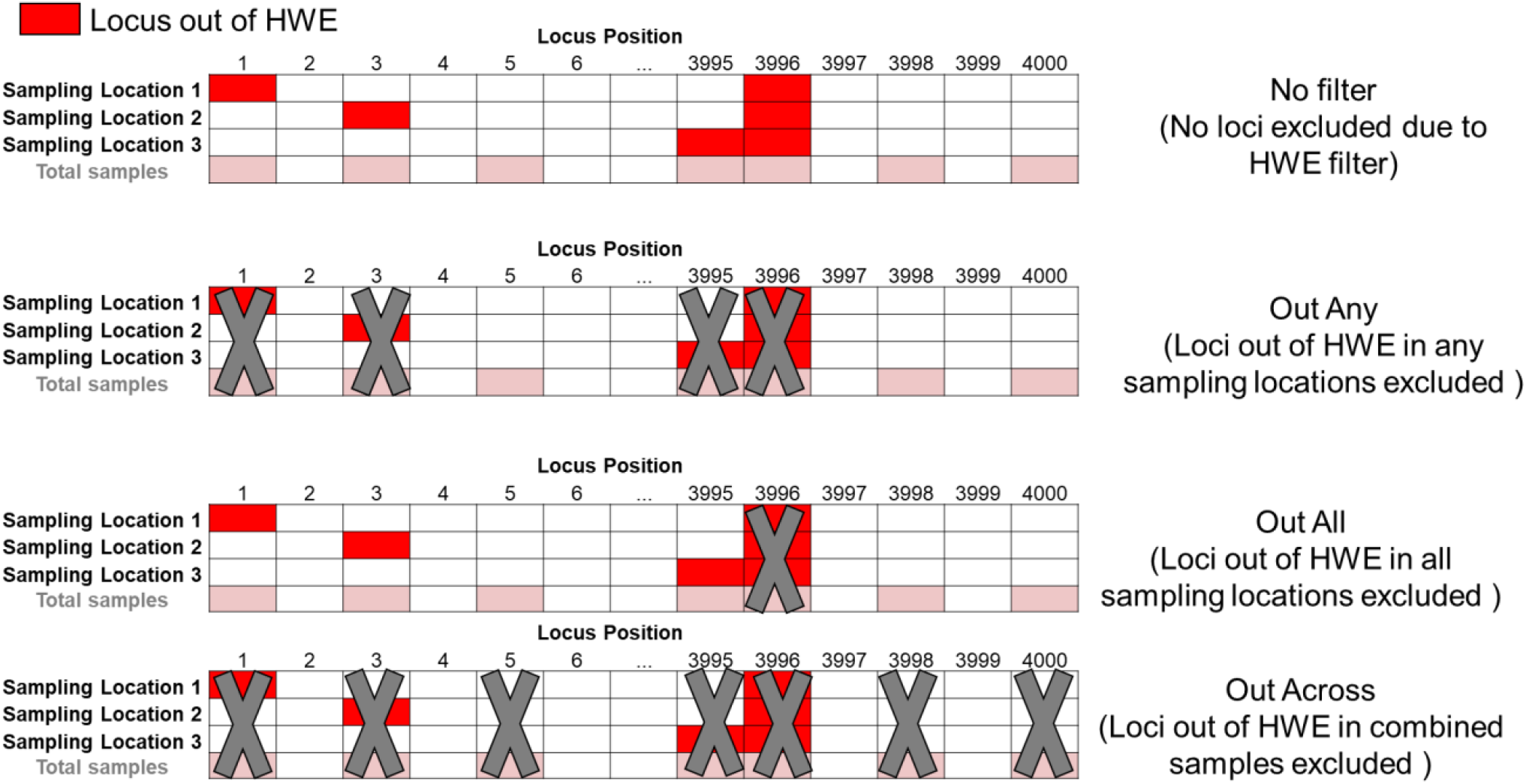
Four commonly applied Hardy-Weinberg Equilibrium (HWE) filtering options (loci removed indicated by grey crosses). In the case of ‘No Filter’, no loci are removed, even if they exhibit departures from HWE. In the case of ‘Out Any’ and ‘Out All’, loci are removed if they exhibit departures from HWE in either any sampling location, or all sampling locations respectively. ‘Out Some’ can be considered a subset of ‘Out All’, where loci are removed if they are out of HWE in a certain proportion of populations. Finally, in ‘Out Across’, loci are removed if they exhibit HWE departures when sampling locations are grouped together.

The ‘Out Across’ approach removes genetic loci that depart from HWE across the entire genomic dataset. This filtering scheme will have a substantial impact on downstream analyses since loci that are strongly informative for population structure are likely to be removed by this filter due to the differences in allele frequencies between populations leading to these loci to being out of HWE when analysed at the total dataset level. However, applying ‘No Filter’ could lead to the retention of genotyping errors or of genetic loci under selection which might be problematic in downstream analyses. Filtering some loci according to the ‘Out All’ (or ‘Out Some’) approach might therefore be advantageous: Only loci that depart from HWE in all (or some) populations would be removed, i.e. the loci that are most likely to be problematic. The same applies to the ‘Out Any’ approach, which is extremely conservative in that it removes loci that show departures from HWE even in a single population. However, both approaches (‘Out Any’ and ‘Out All’) require knowledge about the underlying population structure in order to correctly define populations for assaying patterns of HWE. In the absence of prior knowledge, studies often assume sampling locations to be a proxy for genetic populations. While this assumption might not be problematic in the case of pronounced population structure, conflating sampling location with genetic populations in the case of subtle population structure could be problematic. This is because the application of HWE filters might inflate divergence estimates between sampling locations if they do not accurately map to the underlying population structure. This inflation may occur if loci that discriminate ‘true’ populations were removed through HWE filters, and loci that discriminate sampling locations were retained. This would erroneously reinforce the *a priori* hypothesis that sampling locations reflect underlying genetic populations. This ‘over-splitting’ of populations can be as problematic in a conservation setting as the previously discussed ‘over-lumping’ of populations (i.e. Wahlund effects) in terms of implementing management recommendations.

Despite the potentially substantial impact of HWE-based filtering approaches, they are frequently misused or their application is not reported at all (Sethuraman et al., 2019). While it has been suggested that HWE filtering is often inadequately described and inappropriately applied (Gruber et al., 2018; Waples, 2015), this has not yet been systematically assessed within the field of RADseq-based population genomic research (Table 1). For example, many widely used filtering tools such as VCFtools (Danecek et al., 2011), plink (Chang et al., 2015), and pegas (Paradis, 2010) calculate HWE departures directly from genetic data rather than utilising a population mapping file. This default behaviour might be desirable when studying a single population, as is often the case in large-scale human genomic studies, but it could be problematic in studies comprising many populations for the reasons outlined above (i.e. the default behaviour would therefore be ‘Out Across’, subject to the impact of the Wahlund effect).

Here, we firstly review the common approaches for HWE filtering currently used in the scientific literature, and then systematically explore the effect of different HWE filtering approaches with the help of simulations and empirical biological datasets across a wide range of realistic levels of population structure. We hypothesise that HWE filtering will have a substantial effect, especially on marginally or non-structured populations. Specifically, we hypothesise that the removal of genetic loci that depart from HWE across populations will reduce estimated population structure, whereas the removal of genetic loci that depart from HWE in any population will increase estimated population structure and divergence by reducing the impact of ‘noisy’ loci resulting from methodological artefacts (e.g. variant calling, null alleles). Finally, we hypothesize that HWE filtering schemes that consider population strata will reinforce the *a priori* sample groupings when genetic populations are conflated with sampling locations.

## Methods

### Literature Review

We conducted a literature review for RADseq-based population genomic research using the Web Of Science (Supplementary Information 1 for specific search terms). From the initial results, we selected studies that contained any of the following terms “Hardy”, “Weinberg”, “HWE” or “Hardy-Weinberg”, and excluded those that met any of the following criteria:

1. Described a new panel of SNPs; these studies mostly describe a very small panel of genetic variants.
2. Studied a single population; studying a single population means that HWE filtering will not have an impact on population structure inference.
3. Focused on human populations; we excluded human datasets to avoid ethical concerns around demarcating human populations and the comparatively rare use of RADseq for humans compared to WGS.
4. Consisted of transcriptome- or RNA-derived genetic variants; these variants are likely to display departures from HWE since they are transcriptionally expressed and therefore more likely to be under selection.
5. Did not explicitly discuss HWE filtering; we were not able to discern if these studies had not applied any filtering or had just not mentioned it. Furthermore, it was difficult to ascertain whether this filter was overlooked or intentionally avoided, and would bring the scope of the literature review beyond what was manageable.
6. Was not based on RADseq data; we focused on RADseq data since allelic dropout can be a substantial source of HWE departures, and RADseq is currently one of the predominant RRS approaches for non-model organism population genetics.

The remaining studies were classified into one of the seven categories described in Table 2 (Note that ‘No Filter’ likely underestimates the number of studies that do not utilize Hardy Weinberg filtering, as studies that do not discuss this would not be included in our search results – as we explicitly search for Hardy Weinberg associated studies).

**Table 2.**
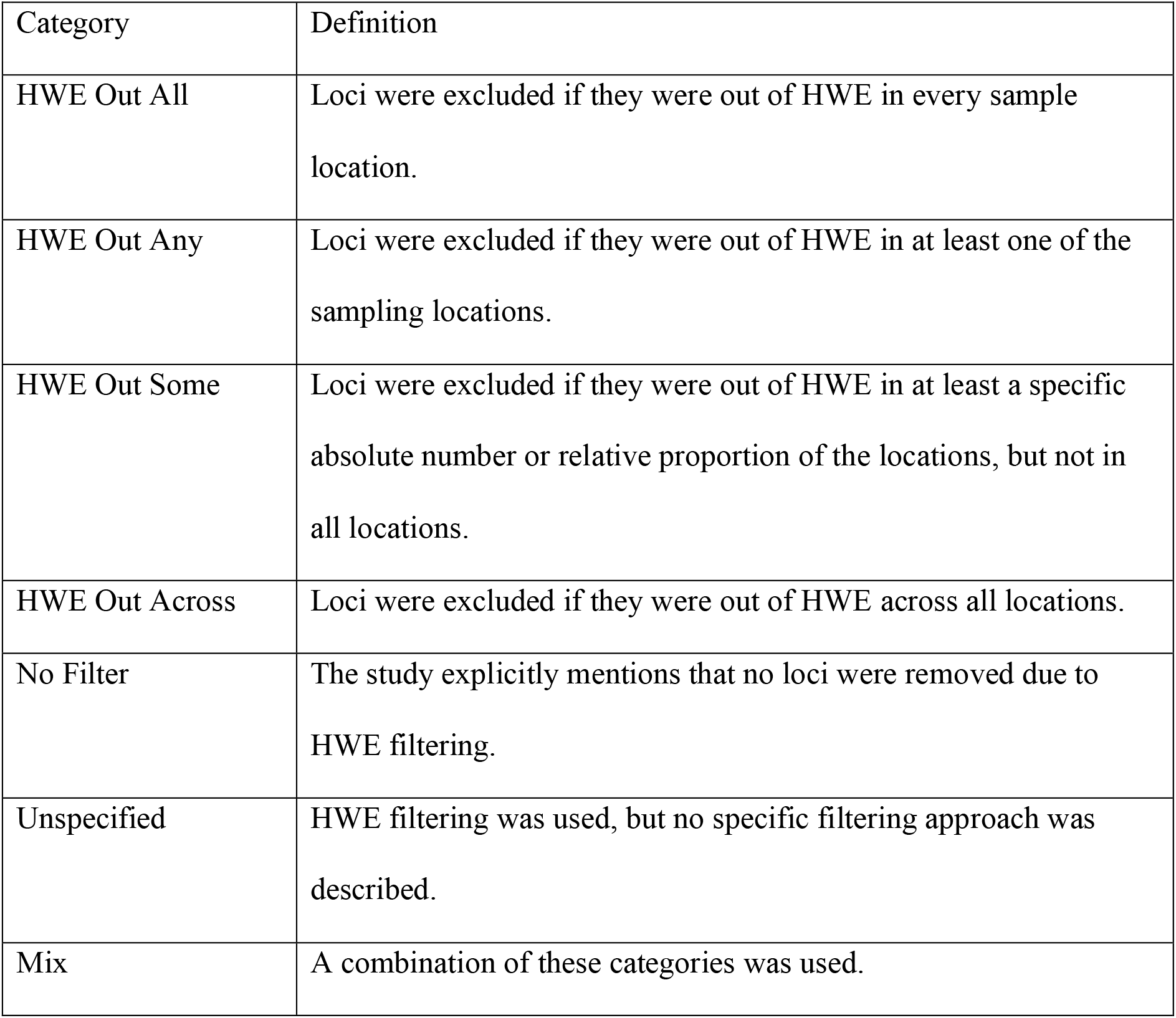
Description of categories used to group scientific studies based on their Hardy Weinberg filtering approaches.

### Simulated data

To investigate the impact of HWE filtering on inference of population structure, we used both simulated and empirical datasets. For all simulations, we used the SLiM forward genetic simulation framework (Messer 2013; Haller and Messer 2017). Due to the availability of well-characterized recombination rates (e.g. Comeron et al. 2012), we simulated a random genome based on the lengths of the 2L, 2R, 3L and 3R chromosomes of *Drosophila melanogaster*. We used the recombination rates determined by Comeron et al. (2012) at 100 kb intervals in combination with the “pseudo-chromosomes” option in SLiM to enable independent simulation of autosomal chromosomes. We assumed a sexually reproducing diploid organism. We chose an arbitrary but realistic mutation rate of 10-8, and an effective population size of 1000. Age-related mortality was implemented with maximum mortality at age seven, with density-dependent survival ensuring fluctuation of the population size around the effective population size.

A single population was created which evolved for 135,000 generations (i.e., three times the number of generations that the initial population took to reach coalescence, namely approximately 45,000 generations), followed by divergence into twelve separate populations with an initial census population size of 80. These populations then evolved for another 15,000 generations with constant migration between adjacent populations (Supp. Fig. 1).

During this period, populations expanded to an effective population size of 1000. Differing migration rates in each scenario adjusted the degree of population structure, with the “Marginal” population structure migration rate at 0.1 (i.e., 0.1 or 10% of a population was transferred to the adjacent population/s in each generation, e.g. population 5 received 10% of both populations 4 and 6), “Low” population structure migration rate at 0.01, “High” population structure migration rate at 0.001, and “Extreme” population structure migration rate at 0.0001. At generation 150,000, 30 individuals were sampled randomly from every other adjacent population, resulting in a total of 180 individuals being sampled from populations 1, 3, 5, 7, 9, and 11 (Supp. Fig. 1).

The resulting VCF was processed by the program RADinitio, which simulates the RADseq process, including restriction enzyme digest and sources of error (e.g., sequencing error, variation in read depth across alleles) (Rivera‐Colón et al., 2021). We used PstI as a restriction enzyme, set mean coverage at 10x, and simulated nine PCR cycles, a read length of 150 bp, and a mean insert length of 350 bp with a standard deviation of 35 bp. The simulated fastq reads were aligned to the reference using BWA v.0.7.17 (Li, 2013; Li & Durbin, 2009); we then used SAMtools v1.10 (Li et al., 2009) to convert the alignments to sorted bam files. SNPs were called using a reference-guided Stacks v2.53 workflow (Rochette et al., 2019). We called Stacks via ref_map.pl using default options: 0.05 as the significance level for calling variant sites (var-alpha) and genotypes (gt-alpha), PCR duplicates were not removed, paired-end reads and read pairing were utilised (i.e., we did not use the rm-pcr-duplicates, ignore-pe-reads, and unpaired flags), the minimum percentage of individuals in a population required to output a locus was zero (--min-samples-per-pop/-r), and the minimum number of populations a locus had to be present in was one (--min-populations/-p). We then used the populations module of Stacks to write one random SNP from each locus to a VCF file as input for downstream analyses (i.e., using the write-random-snp and VCF flags).

### Empirical data

In order to validate our results against empirical data and across multiple SNP calling pipelines, we selected three publicly available datasets as they represented a range of organisms, with a range of population structure: A DArTseq (Diversity Arrays Technology sequencing) dataset of a New Zealand isopod (*Isocladus armatus*) (Pearman et al., 2020), and two RADseq datasets of the New Zealand fur seal (*Arctocephalus forsteri*) (Dussex et al., 2018) and the Plains zebra (*Equus quagga*) (Larison et al., 2021). For the isopod dataset, the DArTseq genotypes were provided by diversityarrays™, who generated them using their proprietary SNP calling software with a *de novo* assembly (SRA: PRJNA643849, https://osf.io/kjxbm/). For the other two datasets, a Stacks workflow similar to the *in silico* analyses was used to generate the SNP genotypes. SRA data (New Zealand fur seal: SRP125920, single-end data; and zebra: SRP288329, paired-end data) was obtained (using prefetch) and converted to fastq (using fastq-dump) with sratoolkit v2.9.6 (Leinonen et al., 2011). Metadata associated with these datasets (Dussex et al., 2018; Larison et al., 2021) was used to generate popmap files. Conspecific genomes were used as references, namely Antarctic fur seal for the New Zealand fur seal analyses (GCA_900642305.1_arcGaz3_genomic: Humble et al., 2018) and horse for the zebra analyses (GCF_002863925.1_EquCab3.0_genomic: Kalbfleisch et al., 2018). The Stacks workflow then followed the previously described workflow for the *in silico* datasets.

### SNP filtering

For both *in silico* and empirical datasets, we filtered data on a minor allele count of 2, missingness of 0.8, and then applied various filtering approaches for SNPs departing from HWE (Fig. 1). SNPs exhibiting departures from HWE corresponding to each filtering scheme (i.e., Out Any, Out All, Out Across) were identified using the function hwe.test in the pegas R package (Paradis, 2010), corrected for multiple testing using a Benjamini-Hochberg correction, and subsequently removed using VCFtools.

### Data analysis

To examine variance in our parameter estimates, we sampled with replacement from the total number of SNPs in the filtered VCF to generate ten VCF files consisting of 4,000 SNPs each. To examine population structure, we conducted Principal Component (PCA), FST, and STRUCTURE analyses. PCAs were conducted in R 4.02 (R Core Team, 2020), using a genotype matrix with scaled genotypes following procedures outlined in Linck and Battey (2019) in the adegenet R package (Jombart & Ahmed, 2011). PCAs were compared using the PC_ST_ metric, which represents one minus the ratio of the mean within-population distance to total-population distance within a PCA. Higher values of PC_ST_ are consistent with higher levels of population structure (see Linck & Battey (2019) for an in-depth explanation). FST was calculated using the R package STaMMP (version 1.6.1) (Pembleton et al., 2013). STRUCTURE was run using an admixture model with no *a priori* information regarding population structure, using a K of 6 for our *in silico* data, or a K equivalent to the number of sampled populations for the real data. Pairwise comparisons of filters within each scenario were tested for significance using Mann-Whitney U tests and Bonferroni adjustment (alpha = 0.05) in R 4.02 using rstatix (version 0.7.0) (Kassambara, 2021; R Core Team, 2020). Figures were created using the tidyverse and cowplot packages (Wickham et al., 2019; Wilke, 2020).

### Randomisations

To examine if filtering could introduce artificial population structure, we took two of the simulated scenarios (Marginal [M=0.1] and Extreme [M=0.0001]) and randomly assigned individuals to populations before repeating the F_ST_ and PC_ST_ analyses. As no population structure would be expected to in these analyses, any increase in observed population structure due to filtering would have been artificially introduced by the respective filtering approach.

## Results

### Literature Review

Our literature review of 219 scientific publications concerning HWE filtering of RADseq data showed that 53.88% of the publications (n=118) specified their HWE filtering approach (Fig. 2A). Overall, 21% of the publications used some intermediate threshold (‘Out Some) to filter SNPs departing from HWE, 10.5% used ‘Out Across’, 10% used ‘Out Any’, 7.8% explicitly chose not to filter for HWE departure and outlined their reasons, and 2.3% used ‘Out All’ (Fig. 2B; see Table 2 for definition of filtering approaches). The remaining 101 publications (46.12% of all publications) did not specify the HWE filtering approach in sufficient detail (Fig. 2A): 45 publications (20.6% of all publications) specified only the filtering tool they used, whereas the remaining publications (25.6% of all publications) did not specify any information (“Unspecified”; Fig. 2C). If the default behaviour of the specified filtering tools is assumed, another 11.9% of all publications (n=26) used ‘Out Across’ (Fig. 2C). Overall, this means that at least 22% of the publications that filtered for departure from HWE have most likely used the ‘Out Across’ approach, but we expect this proportion to be even higher due to the large proportion of unspecified publications. Finally, some publications (8.7%, n=19) used filtering tools that explicitly consider population stratification in HWE calculations (such as Arlequin (Excoffier et al., 2005) or Genepop (Rousset, 2008)), but the publications did not report the exact filtering approach (“Within”, Fig. 2C).

**Figure 2.**
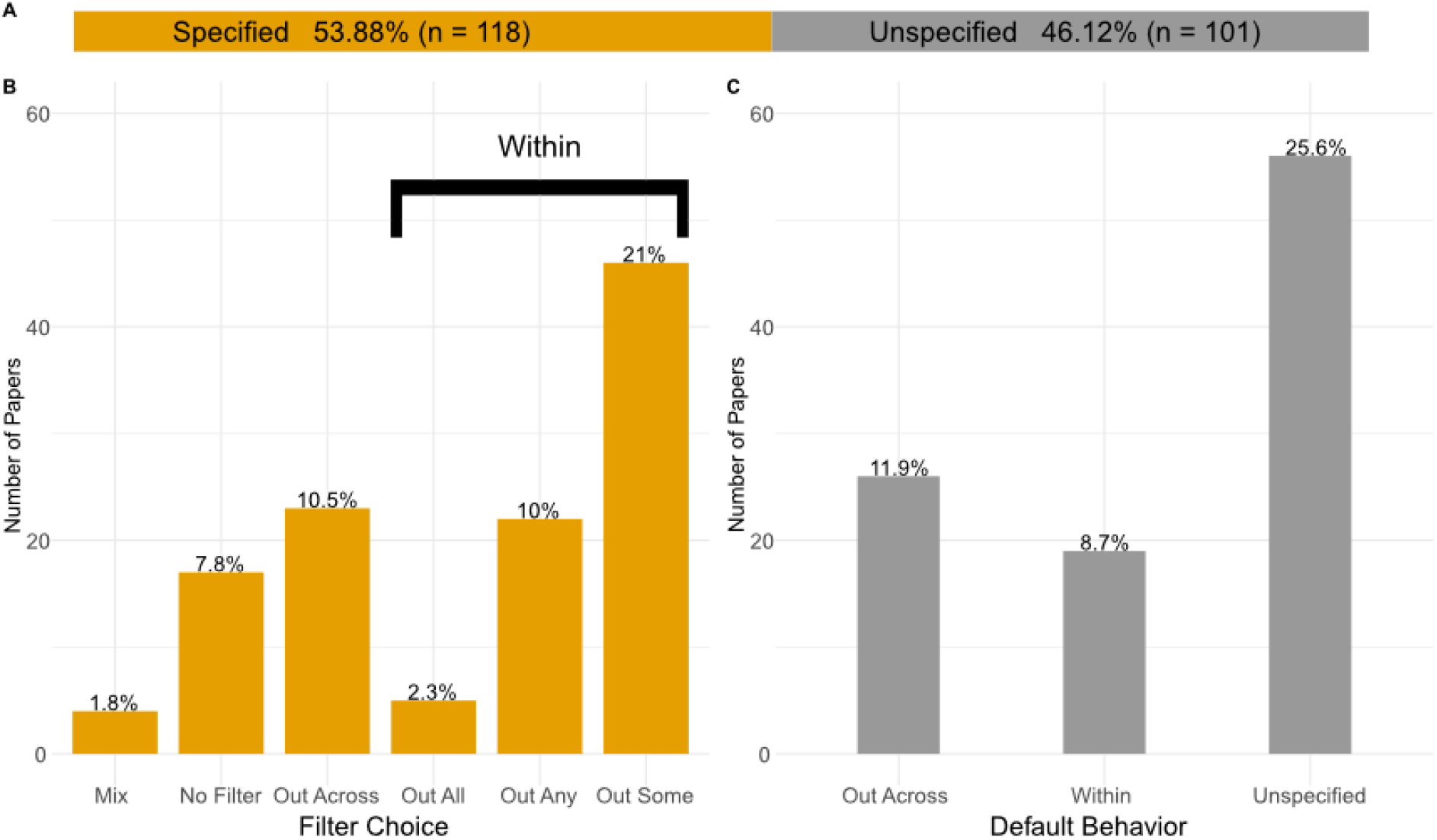
A) Distribution of publications that specified their HWE filtering approach (orange) versus publications that did not specify the approach in sufficient detail (grey). B) The distribution of publications that specified their HWE filtering approach across different filtering schemes: ‘Mix’ (mix of the following filters), ‘No Filter’ (no HWE filter), ‘Out Across’ (loci removed if out of HWE across the pooled dataset), ‘Out All’ (loci removed if out of HWE in each sampling location), ‘Out Any’ (loci removed if out of HWE in any sampling location), and ‘Out Some’ (loci removed if out of HWE in at least a certain number/relative proportion of sampling locations, but not in all locations). C) The distribution of publications that did not specify Hardy-Weinberg filtering approach and with the default behaviour of the filtering tools used (where specified) assumed: ‘Out Across’ (as defined above), ‘Within’ (the paper specified that they used population information for HWE filtering, but not specifically whether this was ‘Out All’, ‘Out Any’, or ‘Out Some’) and ‘Unspecified’ (the paper did not specify the tool).

### *In silico* data analysis

Measurements of population stratification extracted from PCAs (PC_ST_) were largely robust across different HWE filtering approaches regardless of population structure, with the exception of ‘Out Across’ (Fig. 3).

**Figure 3.**
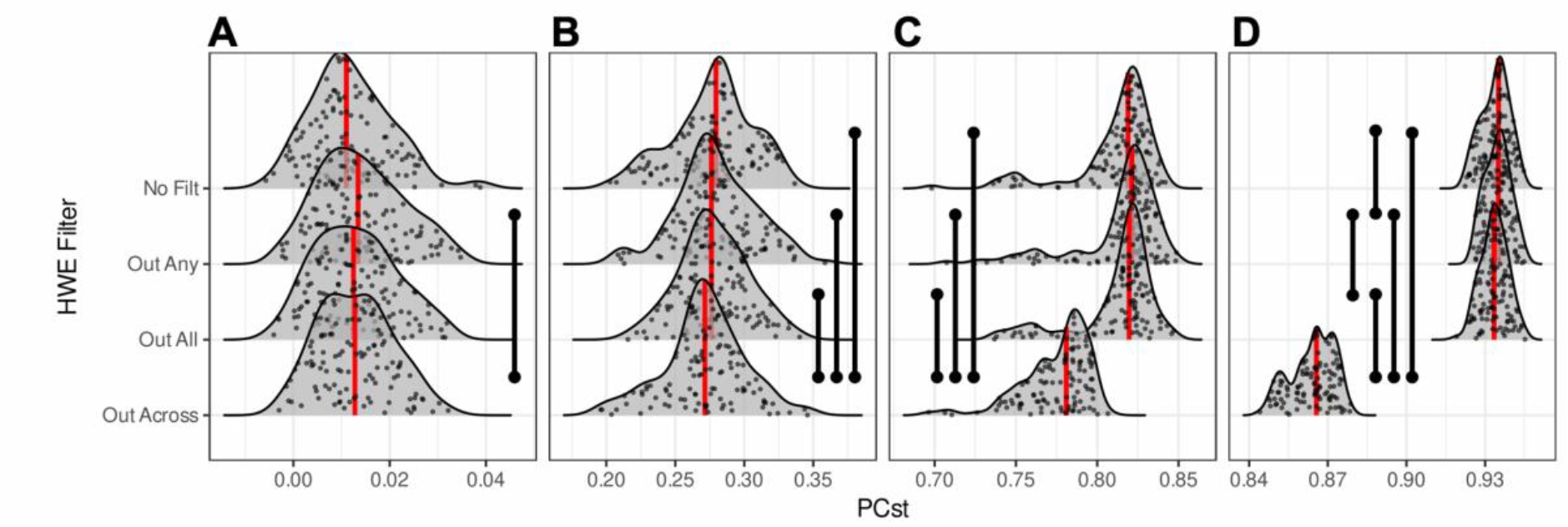
Distributions of PC_ST_ across HWE filtering approaches and degrees of inferred population structure. A represents marginal population structure (i.e. high migration, M=0.1), B represents low population structure (M=0.01), C represents high population structure (M=0.001), and D represents extreme population structure (i.e. low migration, M=0.0001). Red lines indicate median values, black vertical bars indicate statistically significant comparisons (Mann-Whitney U tests, Bonferroni adjustment).

The effect of ‘Out Across’ became apparent with increasing population structure, reducing PC_ST_ estimates in comparison with other filtering approaches (Fig. 3). The remaining filtering approaches delivered qualitatively similar PC_ST_ estimates (except for extreme population structure where all filtering approaches led to different results but ‘Out Across’ still dominated the divergence in PCst estimates; Fig. 3D). This indicates that the ‘Out Across’ filter reduces estimated population structure evident in a PCA in relation to the other filtering schemes.

In the case of F_ST_, we similarly observed an increasingly strong effect of the ‘Out Across’ filtering approach on reducing inferred F_ST_ with increasing levels of population structure (Fig. 4). While ‘Out All’ and ‘No Filter’ consistently delivered similar F_ST_ estimates, we found that ‘Out Any’ led to larger inferred F_ST_ values, with the exception of extreme population structure where F_ST_ was slightly (but significantly) reduced for this filtering approach.

**Figure 4.**
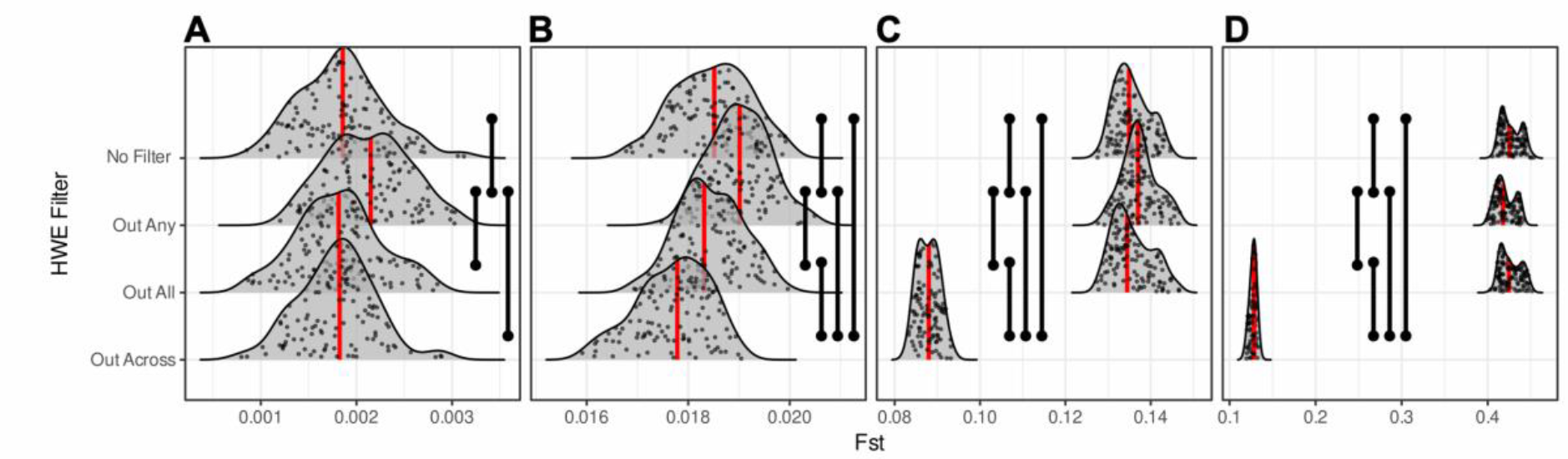
Distributions of inferred F_ST_ across HWE filtering approaches and degrees of inferred population structure. A represents marginal population structure (i.e. high migration, M=0.1), B represents low population structure (M=0.01), C is high population structure (M=0.001), and D represents extreme population structure (i.e. low migration, M=0.0001). Red lines indicate median values, black vertical bars indicate statistically significant comparisons (Mann-Whitney U tests, Bonferroni adjustment).

For the STRUCTURE analyses, we observed that ‘Out Any’ and ‘Out Across’ filters significantly increased the average nucleotide distance between inferred population clusters in the marginal and low population structure scenarios, while ‘Out Across’ decreased the inferred amount of structure in the high and extreme population structure scenarios (Fig. 5).

**Figure 5.**
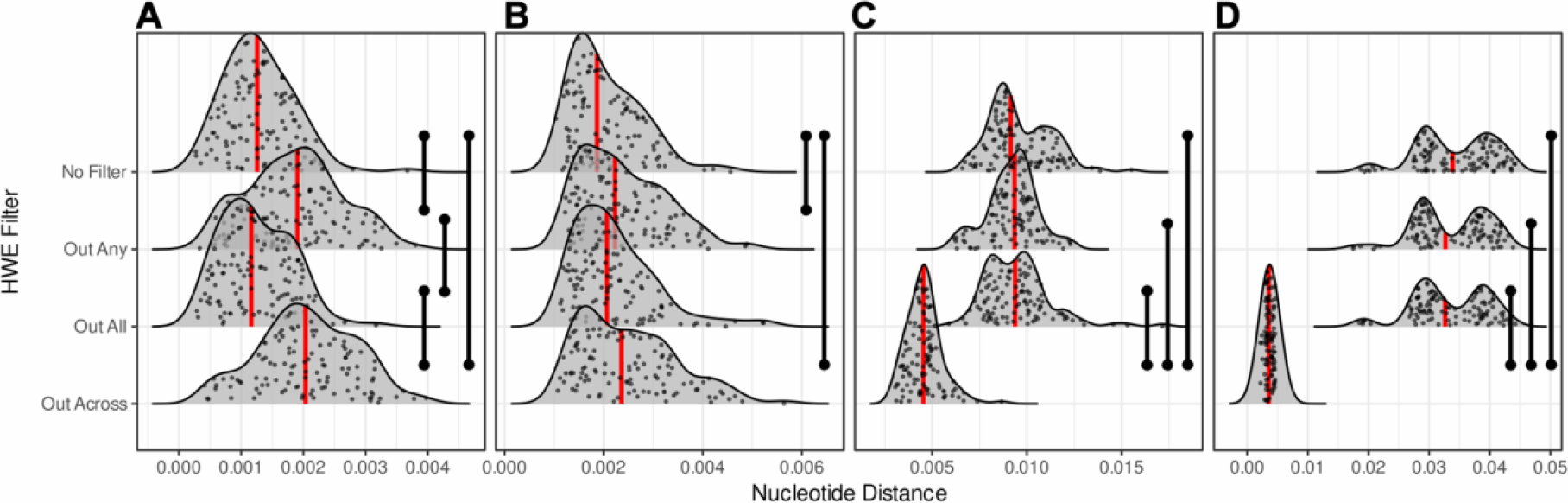
Distributions of average nucleotide distance between inferred population clusters from STRUCTURE, across differing filtering regimes and levels of population structure. A represents marginal population structure (i.e. high migration, M=0.1), B represents low population structure (M=0.01), C is high population structure (M=0.001), and D represents extreme population structure (i.e. low migration, M=0.0001). Red lines indicate median values, black vertical bars indicate statistically significant comparisons (Mann-Whitney U tests, Bonferroni adjustment).

### Randomised data

In the randomized datasets, PC_ST_ distributions were broadly similar across filtering regimes in the case of marginal population structure (Fig. 6A). In the case of extreme population structure scenario (Fig. 6B), the filtering schemes ‘No Filter’, ‘Out Any’ and ‘Out All’ were all significantly different to ‘Out Across’, all leading to slightly higher levels of structure. Given, however, that the ‘No Filter’ approach led to significantly higher estimated structure than the ‘Out Across’ approach, this suggests that our filtering approaches do not lead to any spurious inference of structure for panmictic scenarios. Similar results were obtained for F_ST_ estimates (Fig. 7).

**Figure 6.**
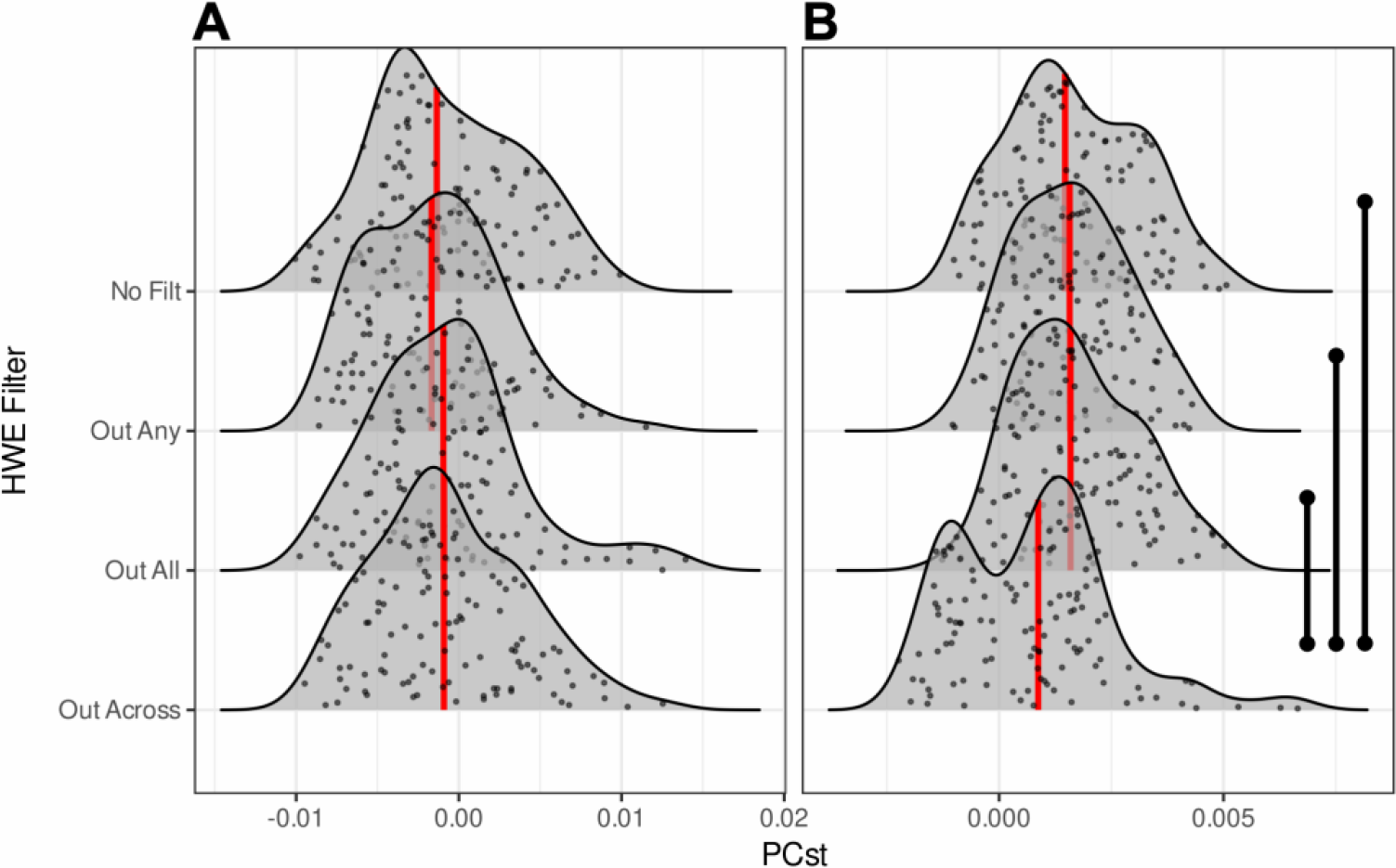
Distributions of PC_ST_ of the randomized SNP datasets across HWE filtering approaches. A represents marginal population structure (A; i.e. high migration M=0.1) and B represents extreme (M=0.0001) population structure. Red lines indicate median values, black vertical bars indicate statistically significant comparisons (Mann-Whitney U tests, Bonferroni adjustment).

**Figure 7.**
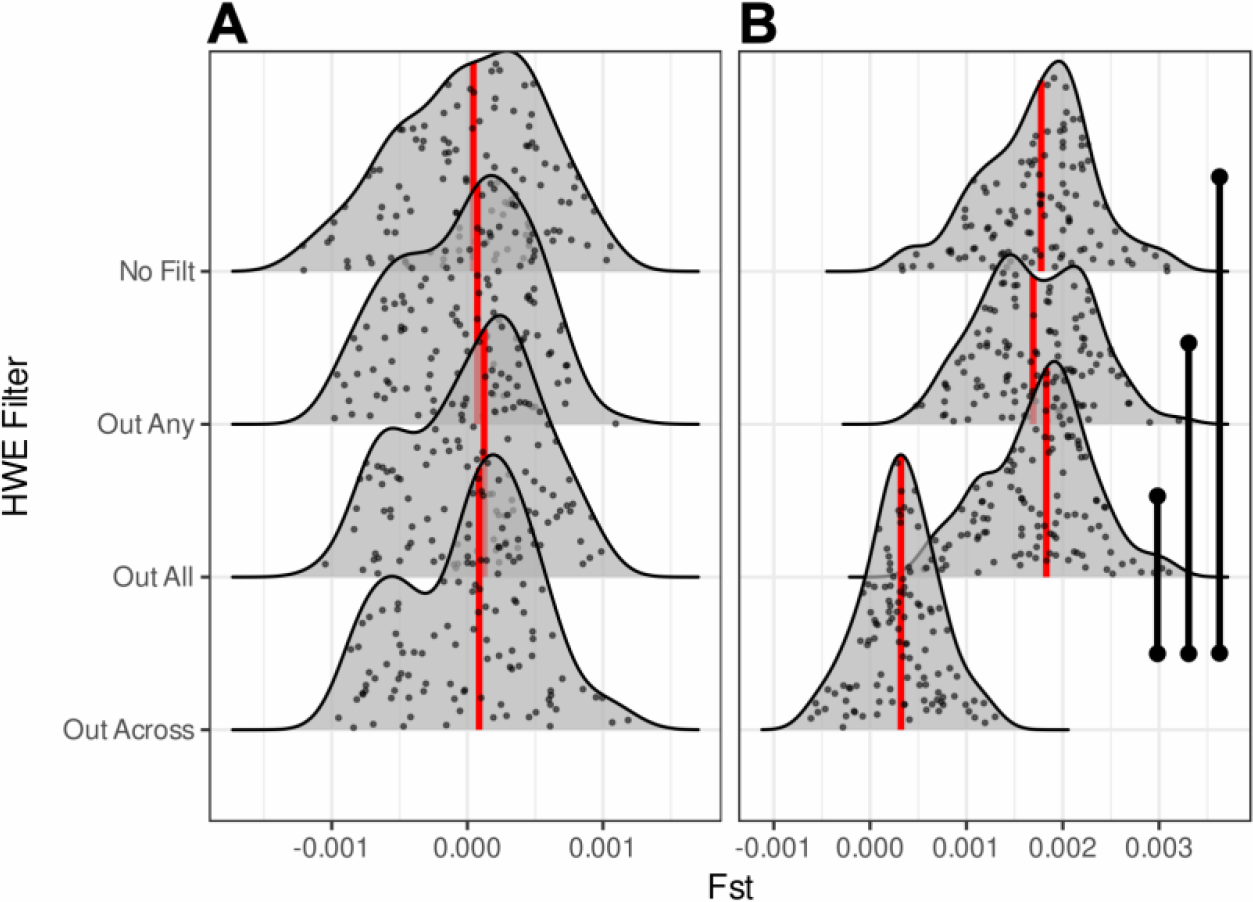
Distributions of F_ST_ of the randomized SNP datasets across HWE filtering approaches. A represents marginal population structure (A; i.e. high migration M=0.1) and B represents extreme (M=0.0001) population structure. Red lines indicate median values, black vertical bars indicate statistically significant comparisons (Mann-Whitney U tests, Bonferroni adjustment).

### Empirical data analysis

The results from the empirical datasets were generally concordant with those from the simulations. No significant differences were observed among filters for PC_ST_ in the species with the weakest population structure, the New Zealand fur seal (Fig. 8A). In the species with more pronounced population structure (zebra and isopod, Fig. 8B-C), the ‘Out Across’ filter had significantly reduced PC_ST_ in comparison with the other filters. ‘Out Any’ had marginally higher estimated structure than ‘No Filter’ or ‘Out All’ in the isopod dataset.

**Figure 8.**
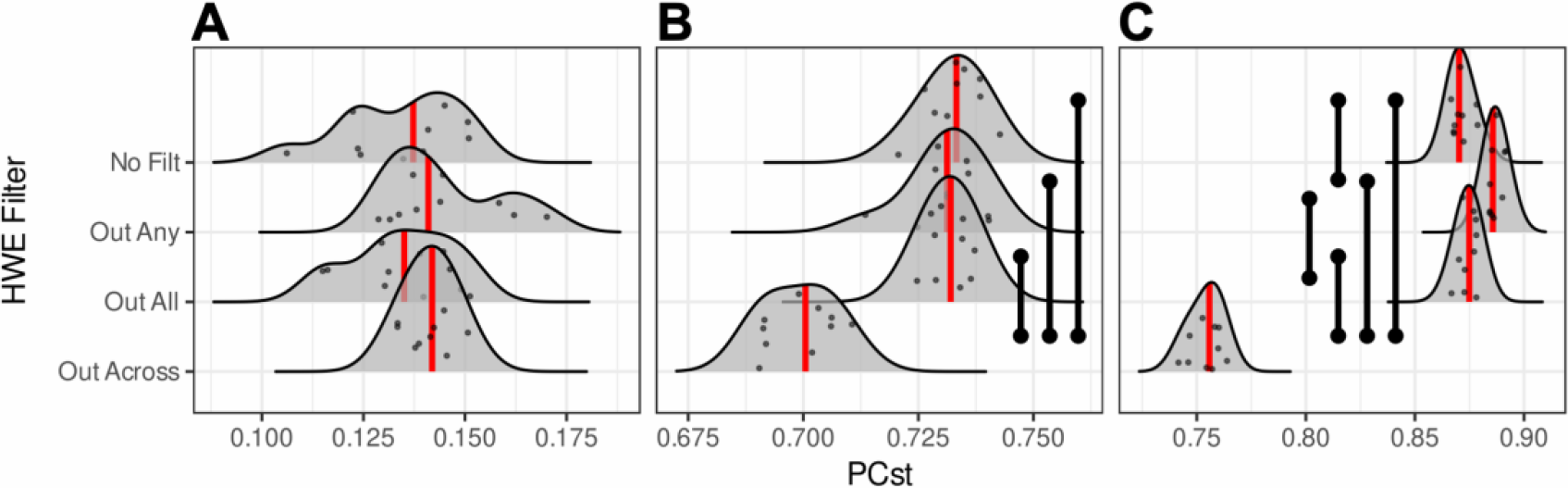
PC_ST_ distributions for empirical datasets, A represents New Zealand fur seal data (*Arctocephalus forsteri*), B represents from the Plains zebra (*Equus quagga*), and C represents a New Zealand isopod (*Isocladus armatus*). Red lines indicate the median value for each distribution, black vertical bars indicate statistically significant comparisons (Mann-Whitney U tests, Bonferroni adjustment). Species ordered from low population structure (New Zealand fur seal) to high population structure (isopod).

Similar results were obtained for F_ST_ (Fig. 9), where the filtering approaches had only small impacts for the inference of population structure in the species with low population structure (New Zealand fur seal), while ‘Out Across’ significantly reduced F_ST_ estimates for the species with higher levels of population structure (Plains zebra and isopod).

**Figure 9.**
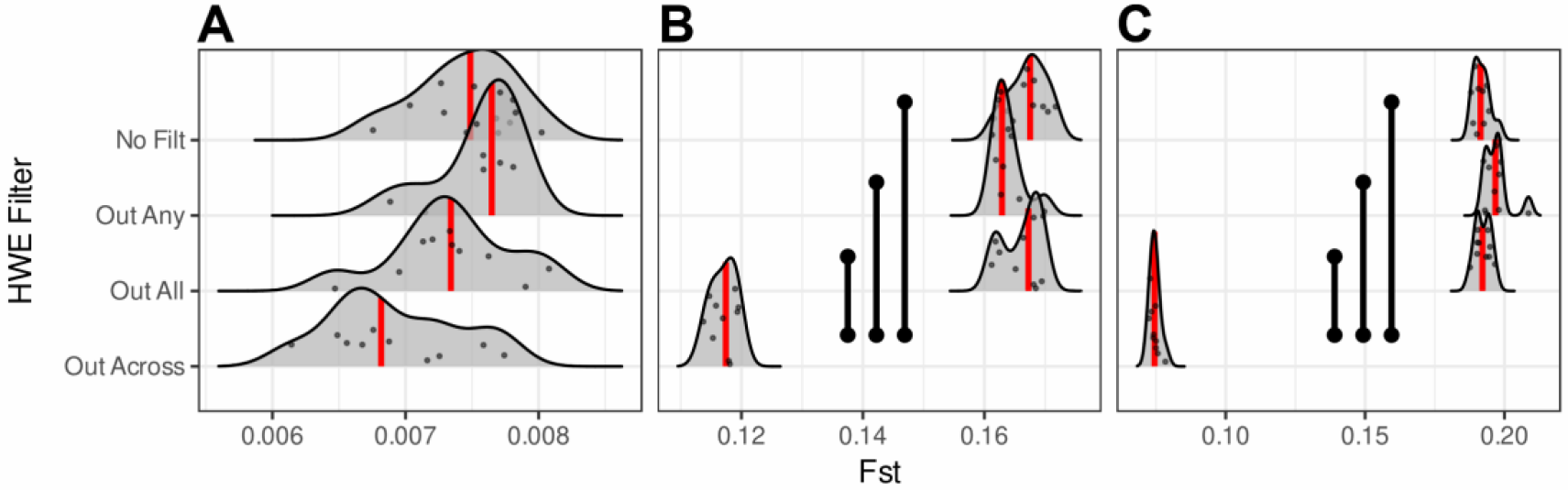
F_ST_ distributions for empirical datasets, A represents New Zealand fur seal data (*Arctocephalus forsteri*), B represents from the Plains zebra (*Equus quagga*), and C represents a New Zealand isopod (*Isocladus armatus*). Red lines indicate the median value for each distribution, black vertical bars indicate statistically significant comparisons (Mann-Whitney U tests, Bonferroni adjustment). Species ordered from low population structure (New Zealand fur seal) to high population structure (isopod).

The ‘Out Across’ filtering approach similarly reduced the estimated nucleotide distance between clusters for zebra and isopod (the species with the most marked population structure). In addition, the ‘Out Any’ filtering approach led to a significant reduction in estimated nucleotide distance in the isopod dataset (Fig. 10).

**Figure 10.**
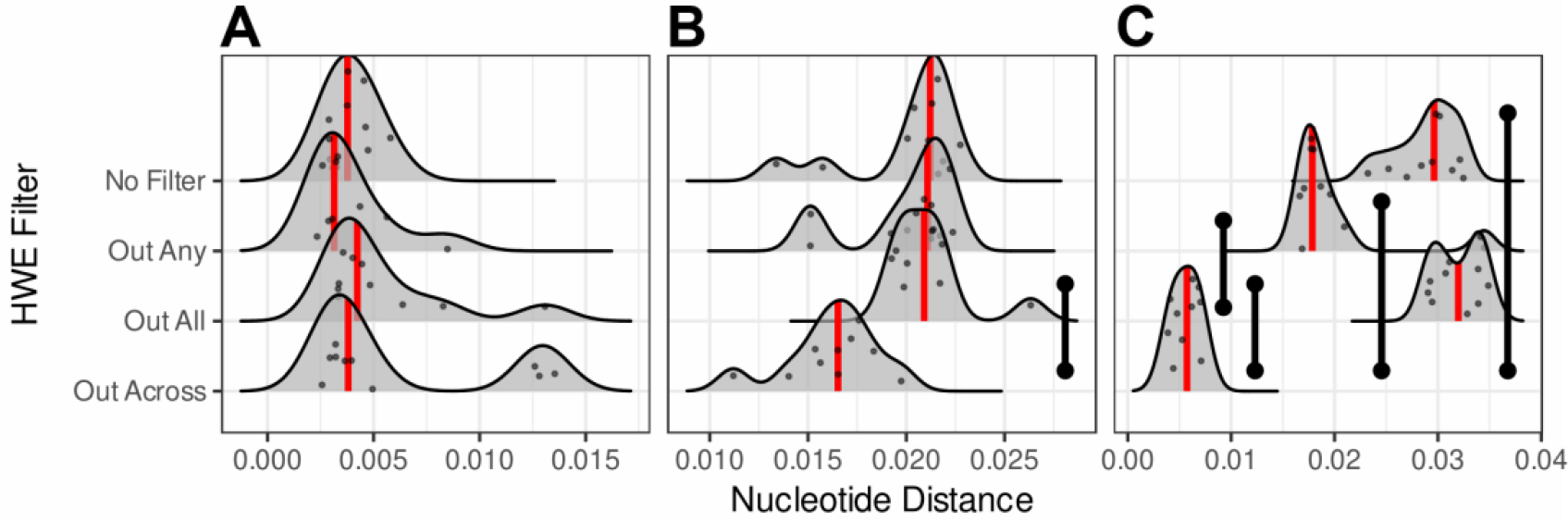
Nucleotide distance distributions for empirical datasets, A represents New Zealand fur seal data (*Arctocephalus forsteri*), B represents from the Plains zebra (*Equus quagga*), and C represents a New Zealand isopod (*Isocladus armatus*). Red lines indicate the median value for each distribution, black vertical bars indicate statistically significant comparisons (Mann-Whitney U tests, Bonferroni adjustment). Species ordered from low population structure (New Zealand fur seal) to high population structure (isopod).

## Discussion

There are many good reasons to impose a filter for HWE, such as removal of loci under extreme selection, paralogs, and sequencing or library preparation artifacts. Thus, HWE filtering can be helpful in standardizing and denoising a dataset. However, in this paper, using both empirical and simulated datasets, we demonstrate that filtering SNPs based on HWE can have substantial impacts on population genetic inferences. In particular, we found that the ‘Out Across’ filtering approach, where loci that depart from HWE across all pooled samples are removed, significantly reduces the amount of inferred population structure relative to ‘No Filter’ or other filtering approaches. This occurs because this filter leads to the inadvertent introduction of a Wahlund effect by not considering any existing population structure, with loci important for delineating population structure being removed by the HWE filter. Despite the strong impact of HWE filtering, our literature review shows that the vast majority of scientific publications that report filtering for HWE do not include sufficient detail to allow replication of this aspect of their analyses. This often occurs because only the filtering tool or significance threshold is reported, while population stratification for filtering is not defined. When default behaviour of filtering tools is assumed, up to 50% of publications may be misapplying HWE filtering (Fig. 2), by using the ‘Out Across’ filtering approach. Some commonly used filtering tools such as VCFtools and plink do not consider population structure when calculating deviations from HWE, and therefore the reliance on default settings may lead to the removal of the very loci that are informative for population structure. Importantly, even the implementation of an extremely conservative significance level for identifying “problematic” loci will not solve the issues of the ‘Out Across’ filtering approach, as an extreme Wahlund effect will be observed in instances of extreme population structure – which would naturally draw loci closer to even stringent significance levels.

We hypothesized that 1) use of an ‘Out Across’ filter would substantially reduce inferred population structure, and 2) that the use of an ‘Out Any’ filter would lead to an increase in inferred population structure. Consistent with these hypotheses we found that 1) filtering across populations (‘Out Across’) had the greatest effect, substantially reducing inferred population structure, and 2) filtering loci that were out of HWE in any population (‘Out Any’) had a marginal, but consistent effect in increasing the degree of estimated population structure in the case of F_ST_ inference (but not in the cases of STRUCTURE or PC_ST_ analyses).

### Impact of filtering on different measures of population structure

PC_ST_ is a non-parametric measure of population structure developed by Linck and Battey (2019) to standardize comparisons of PCAs. In contrast, F_ST_ and nucleotide distance (inferred from STRUCTURE) are widely used parametric analyses that have explicit underlying biological assumptions.

Contrary to our hypothesis where we assumed the ‘Out Any’ filter would strengthen the inference of population structure due to the removal of ‘noisy’ loci, we observed little to no effect of this filter on PC_ST_ in any of our simulations. The lack of effect of ‘Out Any’ on PC_ST_ may be explained by the fact that PCA (1) makes no assumptions about the underlying population structure, (2) is non-parametric, or (3) that PC_ST_ is calculated based on only the first ten principal components, thereby limiting the impact of ‘noisy’ loci on this metric due to the first ten principal components capturing only the majority of the variation.

In contrast to the PC_ST_ results, for two different parametric methods – STRUCTURE and F_ST_ – different filtering approaches strongly impacted inferred estimates of population structure. For inferred F_ST_ we observe that, with the exception of the extreme population structure scenario (i.e. low migration [M=0.0001]), ‘Out Any’ tended to lead to inference of marginally higher structure than other filters, in line with our hypothesis that this filter would strengthen inference of population structure. The increase in observed F_ST_ in these scenarios (low population structure [M=0.1] to high population structure [M=0.001]) is indicative that filtering using an ‘Out Any’ approach may increase the ability to detect marginal population structure. This inference of marginal structure does not appear to be artificially introduced due to the filtering regime, as when population allocations are randomized – the filtering regime did not introduce artificial structure (Fig. 7). This is in contrast to our hypothesis that filtering approaches might reinforce the structure between *a priori* groupings corresponding to sampling locations, rather than “true” underlying populations.

Similarly, with the exception of marginal population structure (i.e. high migration [M=0.1]), ‘Out Across’ resulted in reduced inferred population structure in comparison to the other filtering approaches. In the marginal population structure scenario, the migration rate was so high that it is likely that all sampling locations could be considered a single population; therefore, the use of ‘Out Across’ did not have any major impact.

In the case of STRUCTURE analyses, we used the average of the nucleotide distance matrix from the STRUCTURE output as a metric to compare analyses, with larger average nucleotide distances between inferred clusters indicative of greater population structure. We found that at lower levels of underlying population structure, the filtering approaches had a greater impact on STRUCTURE results, with ‘Out Across’ and ‘Out Any’ both leading to marginally higher inferred population structure than the other two filters. As population structure increased, these effects were reduced and ‘Out Any’ became comparable with other filters, while ‘Out Across’ increasingly reduced the average nucleotide distance between populations.

The observation of a reduction in inferred structure associated with filtering across populations (‘Out Across’) can be largely attributed to the introduction of a Wahlund effect, where loci that are informative for population structure (i.e., fixed in one population but not another) are removed due to exhibition of a reduction in heterozygosity as assessed across the total pooled samples. The observation of an increase in inferred population structure associated with filtering loci that depart from HWE in any population (‘Out Any’) could possibly be explained by the selection of loci that conform best to the *a priori* population groupings. However, in our analyses of simulated panmictic populations, we did not find that the ‘Out Any’ filtering approach introduced artificial structure. Instead, we conclude that this filtering approach largely increases estimates of pre-existing structure rather than introducing artificial structure, potentially by removing ‘noisy’ loci that are not consistently found out of HWE in each population, but likely would be found to be out of HWE if per-population sample sizes were larger.

### Comparison to empirical data

Broadly, the patterns observed in our simulated data were also observed, albeit to a slightly lesser extent, in empirical datasets. Specifically, ‘Out Across’ tended to reduce the inferred amount of population structure for the Plains zebra and New Zealand isopod – both of which have generally high population structure in all other analyses, while for the New Zealand fur seal, no effect of ‘Out Across’ was observed – consistent with our observations of low population structure in the simulated datasets. However, some discrepancies were observed – for F_ST_, the Plains zebra dataset showed reduced inferred population structure in the case of the ‘Out Any’ filtering approach – contrasting with an increased F_ST_ in the simulations with comparable population structure. However, this difference was not statistically significantly different from any other filtering approach except ‘Out Across’. We further found a significant reduction in STRUCTURE-inferred average nucleotide distance for the New Zealand isopod when comparing the ‘Out Any’ filter approach with ‘No Filter’ or ‘Out All’, while our comparable simulations showed no effect of this filter on inferred population structure via STRUCTURE. The discrepancies between the simulated and isopod analyses likely arise from the fact that simulations do not encapsulate the full complexity of real populations: Our simulations do not consider selection, while the isopod dataset was based on morphotypes thought to be under selection (Pearman et al., 2020; Wells & Dale, 2018).

#### Conclusions and recommendations

We conclude that, despite being a widely used filtering approach, filtering across populations (‘Out Across’) is inappropriate and leads to reduced levels of inferred population structure – especially when population structure is high. Removing loci exhibiting HWE departures in any population (‘Out Any’) can marginally increase the ability to detect population structure in datasets. The impact of removing loci that exhibit departures in every single population (‘Out All’) is similar to not filtering at all (‘No Filter’). Thus, we suggest that authors conduct thorough exploratory analyses before applying HWE filters, and in general avoid the use of an ‘Out Across’ filter. Instead, the application of either a ‘No Filter’ or ‘Out All’ regime should be considered. While ‘Out Any’ is more likely to detect population structure, authors should consider the trade-off between the number of loci lost through application of this filter relative to the information gained.

## Acknowledgements

We would like to acknowledge the helpful conversations we have had with Olin Silander, Sarah Wells, and members of the Gemmell Lab who helped inform this paper, as well as the helpful contributions of Georgia Tsambos and Ben Haller both of whom helped in the troubleshooting of the simulations. We would also like to acknowledge the authors of the empirical studies who both generated the original datasets, but also made these easily accessible and reproducible. W. Pearman was funded by a University of Otago Doctoral scholarship. L. Urban was funded by an Alexander von Humboldt Research Fellowship and a Revive & Restore Science Catalyst Fund. A. Alexander was funded by a Rutherford Postdoctoral Research Fellowship, Genomics Aotearoa, and the University of Otago. The authors wish to acknowledge the use of New Zealand eScience Infrastructure (NeSI) high performance computing facilities for this research and specifically D. Senanayake for assistance with compute implementation. New Zealand’s national facilities are provided by NeSI and funded jointly by NeSI’s collaborator institutions and through the Ministry of Business, Innovation & Employment’s Research Infrastructure programme. URL https://www.nesi.org.nz.

## Author Contributions

WSP and AA conceived the study. WSP, LU, and AA designed the research and analysed the data. WSP wrote the article with input from both LU and AA.

## Data availability

All R scripts and SLIM scripts are in: https://github.com/wpearman1996/HWE_Simulations References for included datasets are available in the Methods section.

---

**Table 3.**
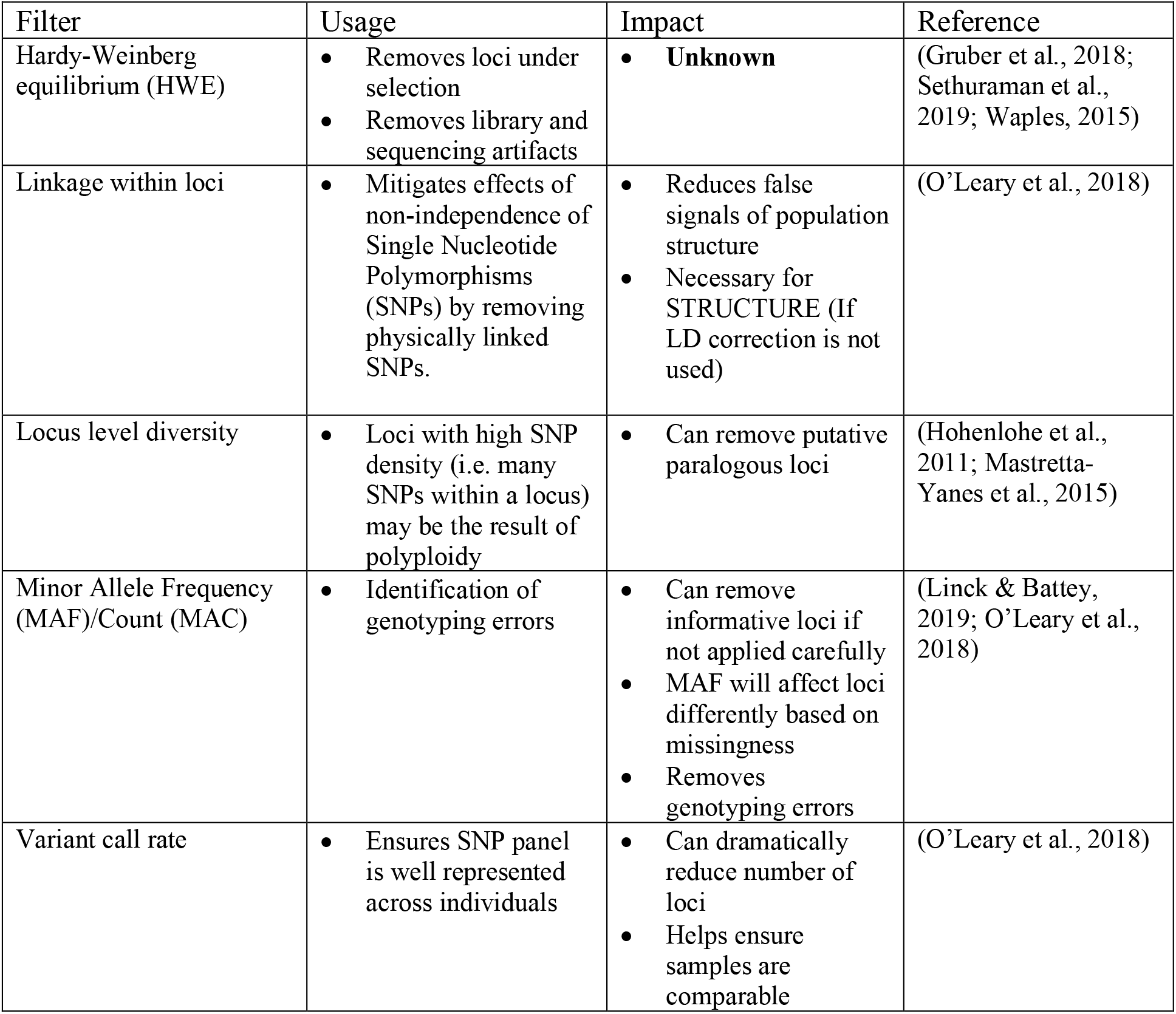
Description of commonly used filtering approaches in the analysis of RADseq data (“Filter”), the reason for their usage (“Usage”), and how they impact population genomic inference (“Impact”).

| Filter | Usage | Impact | Reference |
| --- | --- | --- | --- |
| Hardy-Weinberg equilibrium (HWE) | Removes loci under selection<br>Removes library and sequencing artifacts | **Unknown** | (Gruber et al., 2018; Sethuraman et al., 2019; Waples, 2015) |
| Linkage within loci | Mitigates effects of non-independence of Single Nucleotide Polymorphisms (SNPs) by removing physically linked SNPs. | Reduces false signals of population structure<br>Necessary for STRUCTURE (If LD correction is not used) | (O’Leary et al., 2018) |
| Locus level diversity | Loci with high SNP density (i.e. many SNPs within a locus) may be the result of polyploidy | Can remove putative paralogous loci | (Hohenlohe et al., 2011; Mastretta-Yanes et al., 2015) |
| Minor Allele Frequency (MAF)/Count (MAC) | Identification of genotyping errors | Can remove informative loci if not applied carefully<br>MAF will affect loci differently based on missingness<br>Removes genotyping errors | (Linck & Battey, 2019; O’Leary et al., 2018) |
| Variant call rate | Ensures SNP panel is well represented across individuals | Can dramatically reduce number of loci<br>Helps ensure samples are comparable | (O’Leary et al., 2018) |

**Table 4.** Description of categories used to group scientific studies based on their Hardy Weinberg filtering approaches.

| Category | Definition |
| --- | --- |
| HWE Out All | Loci were excluded if they were out of HWE in every sample location. |
| HWE Out Any | Loci were excluded if they were out of HWE in at least one of the sampling locations. |
| HWE Out Some | Loci were excluded if they were out of HWE in at least a specific absolute number or relative proportion of the locations, but not in all locations. |
| HWE Out Across | Loci were excluded if they were out of HWE across all locations. |
| No Filter | The study explicitly mentions that no loci were removed due to HWE filtering. |
| Unspecified | HWE filtering was used, but no specific filtering approach was described. |
| Mix | A combination of these categories was used. |

