## Supplementary Data for "Commonly used Hardy-Weinberg equilibrium filtering schemes impact population structure inferences using RADseq data"

### Supplementary Material

**Literature search terms**

The following search terms were used in the Web of Science to identify papers of interest:

("population genomics" OR "population genetics" OR "genomics" OR "genetics" OR "genetic" OR "genomic") AND ("SNP" OR "GBS" OR "Genotyping-By-Sequencing" OR "Genotyping By Sequencing" OR "RAD" OR "RADseq" OR "ddRAD" OR "RRS" OR "reduced representation sequencing" OR "reduced-representation sequencing" OR "reduced-representation sequencing" OR "restriction associated" OR "restriction-associated")

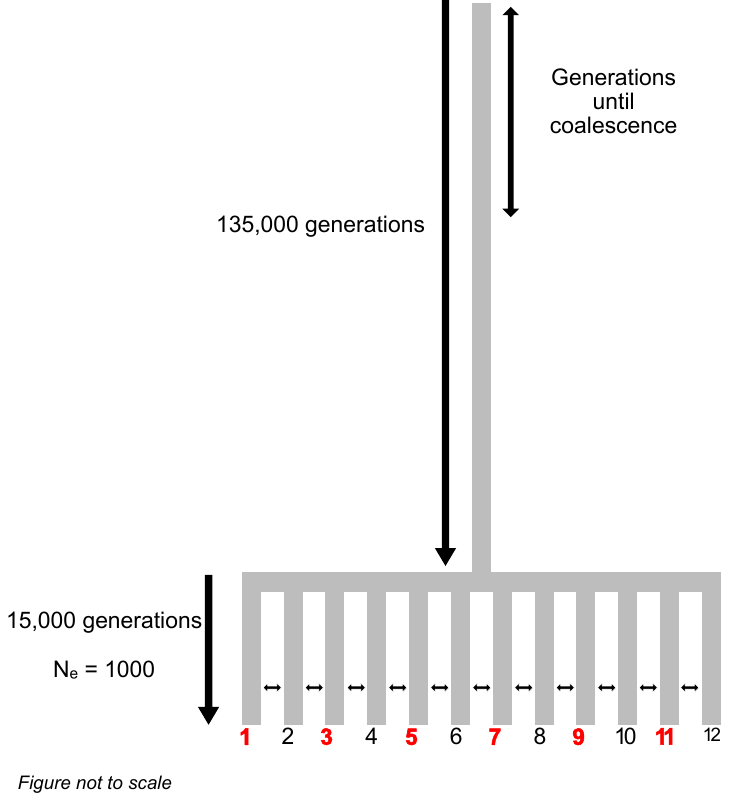

Supplementary Figure 1 Diagram of basic underlying simulation, red labels indicate populations that were ‘sampled’ for genetic data, while double sided arrows indicate two-way migration, which varied from 0.1 to 0.0001.
